# Antibiotic-induced accumulation of lipid II sensitizes bacteria to antimicrobial fatty acids

**DOI:** 10.1101/2022.05.03.490474

**Authors:** Ashelyn E. Sidders, Katarzyna M. Kedziora, Jenna E. Beam, Duyen T. Bui, Joshua B. Parsons, Sarah E. Rowe, Brian P. Conlon

## Abstract

Antibiotic tolerance and antibiotic resistance are the two major obstacles to the efficient and reliable treatment of bacterial infections. Identifying antibiotic adjuvants that sensitize resistant and tolerant bacteria to antibiotic killing may lead to the development of superior treatments with improved outcomes. Vancomycin, a lipid II inhibitor, is of major clinical importance for the treatment of Gram-positive bacterial infections. Here we show that unsaturated fatty acids (UFAs) and vancomycin act synergistically to rapidly kill *S. aureus*, including vancomycin tolerant and resistant populations. Our results suggest that antibiotic-mediated accumulation of lipid II at the septum facilitates membrane invasion by antimicrobial UFAs. UFA-vancomycin dual treatment generates large fluid patches of flexible lipids in the membrane leading to protein delocalization, aberrant septal formation, and loss of membrane integrity. This mechanism of synergy may be exploited for the development of new antibiotic therapies that target lipid II to combat both antibiotic tolerance and resistance.

## Introduction

The 1950’s were the golden age of antibiotic discovery with nearly half of our current arsenal of therapeutics discovered in a single decade^1^. Vancomycin, derived from the word ‘vanquish’, is one of the key discoveries of the golden age that has maintained clinical utility through the present day^2^. Vancomycin is a glycopeptide antibiotic that targets lipid II, an essential membrane-bound building block of the cell wall. Lipid II is synthesized in the cytoplasm and subsequently flipped to the outer leaflet of the cell membrane and incorporated into the mature peptidoglycan layer^3^. Vancomycin binds to the terminal D-Ala-D-Ala residues of lipid II, sterically hindering its incorporation into the growing peptidoglycan layer, resulting in cessation of cell wall biosynthesis and subsequent accumulation of lipid II on the cell surface^4,5^. Since lipid II is highly conserved across all bacterial species, it is often referred to as the “Achilles’ heel” of cell wall biosynthesis and an ideal antibacterial target^6^.

Despite lipid II being highly conserved, resistance to vancomycin is a growing problem and is limiting its utility against some infections^7^. Vancomycin resistance is common and growing in Enterococci. Resistance is mediated by the acquisition of an inducible *van* cassette encoding 7 proteins that generate an alternative lipid II peptide terminus, D-Ala-D-Lac, that significantly reduces vancomycin binding^7^. Such high level vancomycin resistance is also possible in *S. aureus*, although only 14 such cases have been identified to date in the United States^8^. However, vancomycin-intermediate resistant *S. aureus* (VISA), strains which exhibit reduced susceptibility to vancomycin through a variety of non-specific adaptations including a thickening of the cell wall are increasingly common^8^. It is also important to acknowledge that vancomycin treatment frequently fails in the absence of resistance and this failure is frequently attributed to antibiotic tolerance^9^. Antibiotic tolerance is the ability of a bacterial population, or a sub-population of tolerant cells called persisters, to survive high concentrations of bactericidal antibiotics and regrow when antibiotic pressure is removed, leading to re-establishment of the infection^10^. Additionally, antibiotic tolerance can lead to the evolution of resistance^9,11^. Thus, to improve patient outcome we need to develop strategies that can target both antibiotic resistant and antibiotic tolerant populations.

Despite renewed interest in funding antibiotic development, the pipeline is sparse, with few new drugs reaching the market^12^. Developing feasible strategies to revitalize our current arsenal of antibiotics, such as the use of antibiotic adjuvants, which enhance the efficacy of an already approved therapeutic, are highly desirable^13^. Membrane acting agents have frequently been overlooked in drug discovery, as they often lack selectivity for bacterial membranes and display promiscuous activity towards mammalian cells^14^. However, the clinical success and specificity of approved membrane active antibiotics such as daptomycin, polymyxin, and second-generation glycopeptides has highlighted their utility^12^. Additionally, there are several membrane-targeting antibiotic candidates in pre-clinical development including teixobactin, bithionol, and a few antimicrobial peptides^12,15,16^. Our group and others have also previously identified membrane-acting adjuvants that significantly improved the efficacy of aminoglycosides against antibiotic tolerant *S. aureus*^17-21^.

Second-generation glycopeptides, known as lipoglycopeptides, have lipophilic side-chains that interact with the bacterial membrane and display potent bactericidal activity^6^. We were interested in examining the ability of a variety of membrane-acting agents to potentiate vancomycin activity. Here, we identify two antimicrobial fatty acids that are potent vancomycin adjuvants against tolerant and resistant Gram-positive bacteria. Further, we describe the mechanism by which these fatty acids and vancomycin work together to compromise the bacterial cell membrane, leading to rapid cell death.

## Results

### Unsaturated fatty acids potentiate vancomycin killing of *S. aureus*

We have previously shown that rhamnolipids, a biosurfactant produced by *Pseudomonas aeruginosa*, synergizes with aminoglycosides, highlighting the potential of targeting the membrane to enhance antibiotic efficacy against *S. aureus*^18^. We reasoned that cell membrane-acting agents (CMAAs) may also enhance the killing activity of glycopeptides since second-generation glycopeptides have a lipophilic tail and display potent bactericidal activity^22^. We investigated the capacity of 7 CMAAs (Supplementary Table 1) to enhance the bactericidal activity of vancomycin (Fig. 1a). Sublethal concentrations of each CMAA were established experimentally or from our previously published work (Supplementary Figure 1)^18^. Vancomycin and CMAAs have potent bactericidal activity against low density bacterial populations (10^5^ to 10^6^ CFU/ml), but bactericidal activity wanes as population density increases^23-26^. We examined bactericidal activity of vancomycin and CMAAs alone or in combination against *S. aureus* population of 10^8^ CFU/ml, similar to that encounteres in an abscess or infected wound^27-29^. At this density, vancomycin monotherapy resulted in minimal killing (Fig. 1a). Benzyl alcohol, a general membrane fluidizer known to lack antimicrobial activity alone, did not display synergy with vancomycin (Fig. 1a)^25,30^. Lauric acid, a saturated fatty acid, and its monoglyceride derivative, glycerol monolaurate, which synergizes with other antibiotics such as aminoglycosides, did not synergize with vancomycin (Fig. 1a)^8,31-33^. Adarotene, a synthetic retinoid which potentiates aminoglycosides, did not potentiate vancomycin killing (Fig. 1a)^19^. In contrast, rhamnolipids improved vancomycin killing (Fig. 1a), while palmitoleic acid and linoleic acid, two host-produced unsaturated fatty acids, were the most potent adjuvants tested, resulting in a 6-log decrease in viable bacteria after 6 hours (Fig. 1a). Palmitoleic and linoleic acid are both *cis* UFAs, a 16-carbon monounsaturated and an 18-carbon polyunsaturated fatty acid, respectively^25^. Both UFAs have well-established antimicrobial activity and contribute to innate immunity in human nasal secretions, sebaceous glands of the skin, and breast milk ^34,35^. At the bacterial density examined here, neither palmitoleic acid nor linoleic acid had any activity against *S. aureus* in the absence of vancomycin (Fig. 1 and Supplementary Figure 1). However, when these fatty acids were combined with vancomycin we observed over 99% killing of both methicillin-sensitive *S. aureus* (MSSA) and resistant (MRSA) populations after only 30 minutes (Fig. 1b-c). Importantly, the dual treatment eradicated the antibiotic tolerant persister populations as indicated by the absence of a characteristic persister plateau^36^ (Fig. 1b-c).

**Figure 1.**
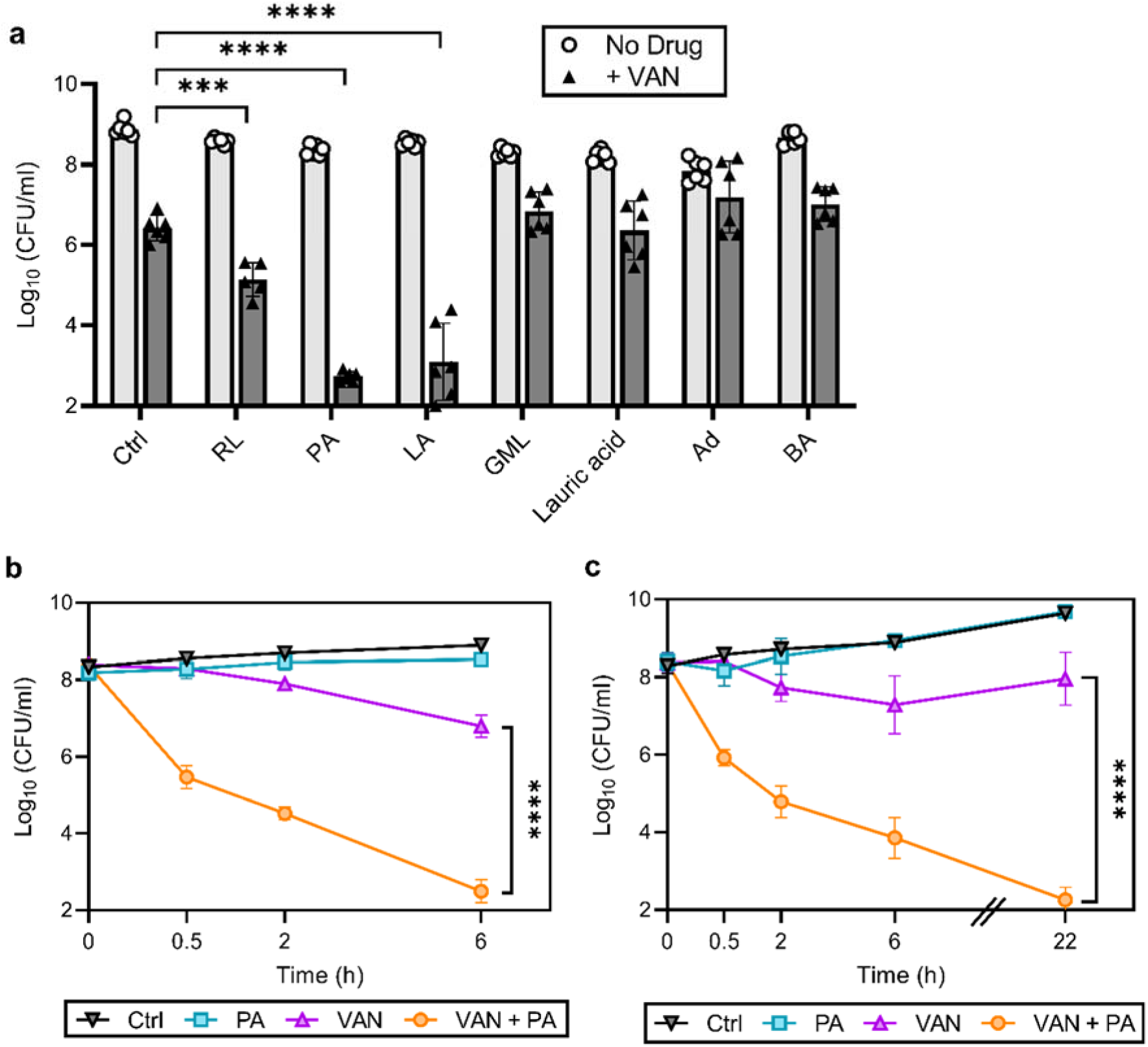
Palmitoleic acid rapidly potentiates vancomycin killing of *S. aureus*. Evaluation of vancomycin (VAN) adjuvant candidates against *S. aureus*. **a** HG003 cultures were grown to exponential phase followed by the addition of sublethal concentrations of CMAAs ± VAN (20 μg/ml, 20X MIC of HG003). Colony forming units (CFU) were enumerated following 6h of treatment. CMAAs include rhamnolipids (RL, 30 μg/ml), palmitoleic acid (PA, 11 μg/ml), linoleic acid (LA, 12 μg/ml), glycerol monolaurate (GML, 30 μg/ml), lauric acid (30 μg/ml), adarotene (Ad, 3.2 μg/ml), and benzyl alcohol (BA, 40 mM). **b**,**c** Survival of HG003 (**b**) or communityacquired MRSA strain LAC (**c**) treated with DMSO (Ctrl, black triangle), PA monotherapy (11 μg/ml, aqua square), VAN monotherapy (20 μg/ml, purple triangle), or PA + VAN combination therapy (orange circle). CFU were enumerated at indicated time points. Data represent the mean values from *n* = 6 biologically independent replicates ± SD. Statistical significance was determined by one-way ANOVA (a) or two-tailed unpaired Student’s *t*-test with a 95% confidence interval at 6h post-treatment (b,c); conditions with significance are indicated on the graph, \*\*\*\**p*<0.0001, \*\*\**p*=0.0003, otherwise comparisons were not significant (n.s.).

### Accumulation of lipid II is necessary for palmitoleic acid potentiation of vancomycin

Because palmitoleic acid exhibited the greatest synergy with vancomycin, we next evaluated whether palmitoleic acid synergizes with other cell wall inhibitors. Antibiotics that target cell wall biosynthesis can be divided into three categories based on their target: early cytoplasmic steps (fosfomycin), membrane bound steps (vancomycin and bacitracin), and assembly/incorporation steps of peptidoglycan (β-lactams)^3^. The latter two categories make up the majority of clinically relevant antibiotics that target the cell wall^3^. To gain insight into the potential mechanism of palmitoleic acid potentiation within the context of the peptidoglycan biosynthesis pathway, we evaluated representative cell-wall acting antibiotics from each category in combination with palmitoleic acid (Fig. 2a). Neither Fosfomycin nor β-lactams exhibited synergy with palmitoleic acid (Fig. 2b-d). Fosfomycin inhibits the cytosolic steps of peptidoglycan biosynthesis by targeting and irreversibly inhibiting enzymes involved in synthesis of peptidoglycan precursors^37^. β-lactam antibiotics, including oxacillin and nafcillin, target penicillin binding proteins (PBPs) which are responsible for incorporating lipid II into the mature peptidoglycan and cross-linking of peptidoglycan^38^. This data indicates that UFAs do not synergize with antibiotics that target the early cytoplasmic steps or the assembly/incorporation steps of peptidoglycan synthesis.

**Figure 2.**
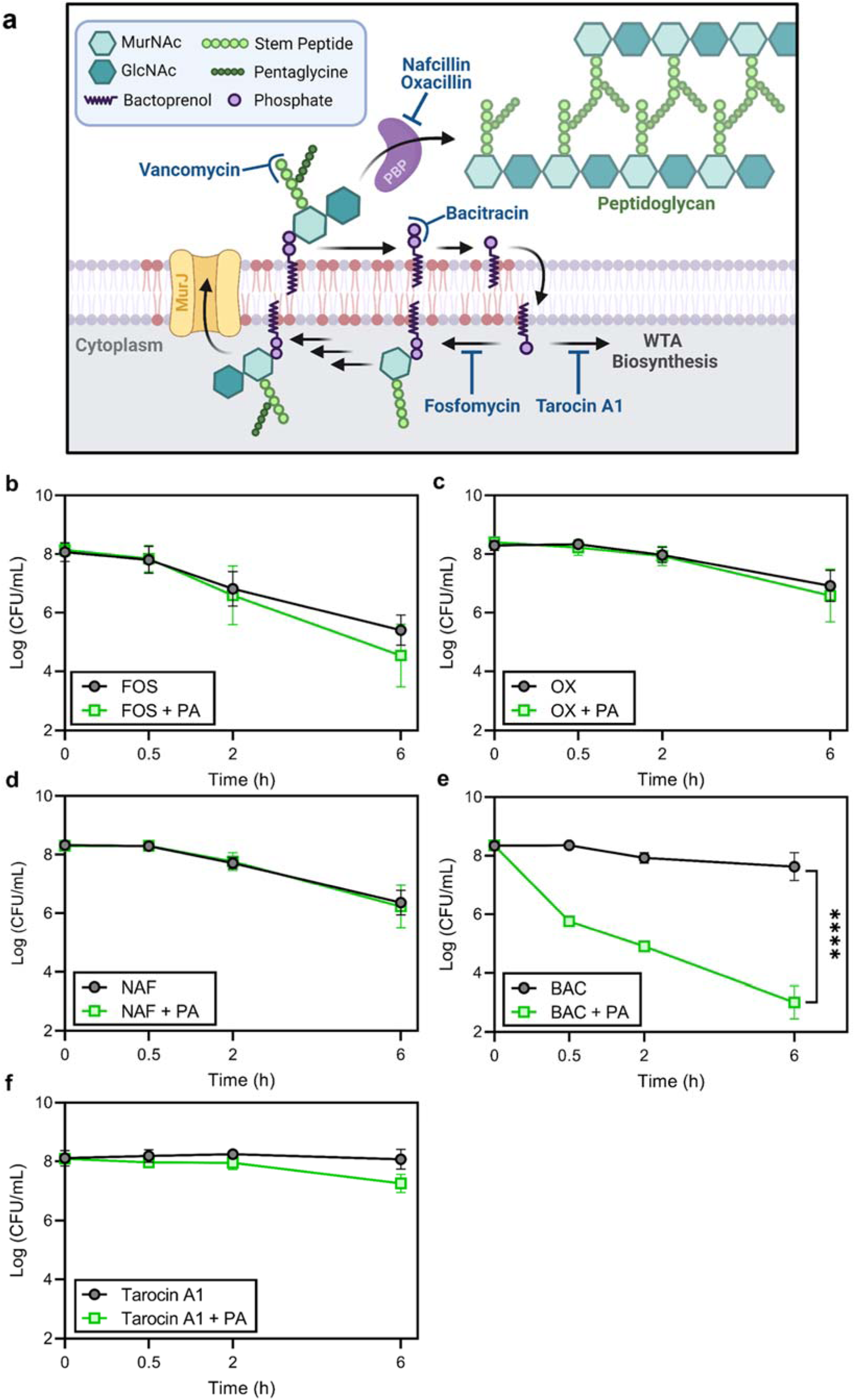
Accumulation of bactoprenol is necessary for palmitoleic acid potentiation of antibiotics. **a** Schematic illustrating the known targets of antibiotics that target the cell wall biosynthetic cycle in *S. aureus*. **b-f** Survival of HG003 challenged with indicated antibiotic monotherapy (black circles) or the antibiotic combined with PA (11 μg/ml, green square). CFU were enumerated at indicated time points. **b** *S. aureus* challenged with fosfomycin (250 μg/ml) ± PA. **c** *S. aureus* challenged with the PBP2 inhibitor, oxacillin (OX, 5X MIC, 5 μg/ml) ± PA. **d** *S. aureus* challenged with the PBP1-4 inhibitor, nafcillin (NAF, 5X MIC, 2.5 μg/ml) ± PA. **e** *S. aureus* challenged with bacitracin (BAC, 250 μg/ml) ± PA. **f** *S. aureus* challenged with tarocin A1 (8 μg/ml) ± PA. Data represent the mean values from *n* = 6 biologically independent replicates ± SD. Statistical analysis was evaluated at the end point by a two-tailed unpaired Student’s *t*-test with a 95% confidence interval; conditions with significance are indicated on the graph, \*\*\*\**p*<0.0001, otherwise comparisons were not significant (n.s.).

Vancomycin inhibits the first surface exposed step of peptidoglycan biosynthesis by binding to the D-Ala-D-Ala moiety of lipid II, preventing lipid II incorporation and thus causing accumulation of lipid II on the surface of the cell^5^. Maximal accumulation of lipid II by vancomycin was found to occur after only 30 minutes^5^. To examine if this accumulation was contributing to the rapid death in the presence of palmitoleic acid, we examined another antibiotic that targets a membranebound stage of cell wall biosynthesis, bacitracin. Bacitracin inhibits the recycling of the essential bactoprenol carrier lipid (Fig. 2a)^39^. Interestingly, we observed that, while bacitracin monotherapy displayed negligible activity, palmitoleic acid potentiated bacitracin killing of *S. aureus* similarly to vancomycin (Fig. 2e, 1b-c). Sequestration of the limited pool of bactoprenol carrier lipids by vancomycin and bacitracin reduces wall teichoic acid (WTA) biosynthesis by redirecting any remaining bactoprenol towards peptidoglycan biosynthesis^40^. Because WTAs have been found to protect *S. aureus* from the antimicrobial activity of UFAs^25^, we assessed whether WTA inhibition was contributed to the mechanism of synergy. Using the WTA inhibitor tarocin A1^41^, we found that inhibition of WTA biosynthesis alone does not synergize with palmitoleic acid, suggesting inhibition of WTA biosynthesis does not contribute to the potentiation of vancomycin or bacitracin (Fig. 2f). Taken together, these results suggest that palmitoleic acid synergy relies on antibiotic induced accumulation of the bactoprenol lipid moiety within the bacterial membrane.

### Dual treatment compromises the permeability barrier function of the bacterial membrane

Palmitoleic acid has previously been reported to cause membrane depolarization and increase permeability, albeit at higher concentrations and at lower cell densities than examined here^25,43,44^. Thus we examined if vancomycin and palmitoleic acid dual treatment caused membrane depolarization. Using the fluorescent voltage sensitive probe DiSC_3_(5)^45^, we found that dual treatment with palmitoleic acid and vancomycin, did not affect membrane potential when added at sublethal concentrations (Fig. 3a). At the highest concentration of vancomycin (20x MIC) combined with palmitoleic acid, some depolarization was observed, however, this was negligible when compared to gramicidin, the pore-forming positive control (Fig 3a).

**Figure 3.**
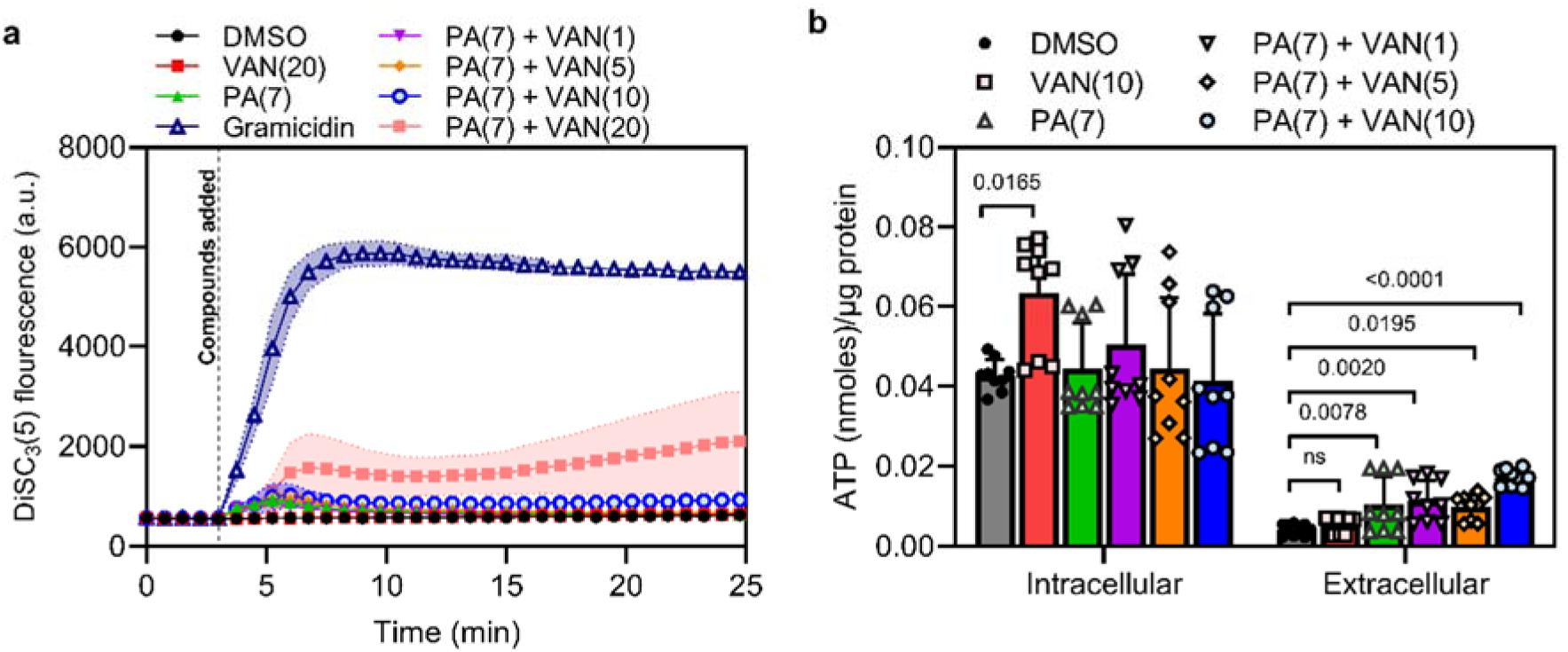
Dual treatment impacts membrane permeability but not energetics. Microtiter plate assays to evaluate respiration and permeability. **a** *S. aureus* grown to late exponential phase was loaded with 7 μM of the voltage-sensitive probe DiSC_3_(5). After the signal stabilized, DMSO, Gramicidin (1 μM), or PA (7 μg/ml) ± VAN (1X, 5X, 10X, or 20X MIC) were added as indicated on the graph. Membrane potential was measured (610/660nm) on a plate reader every 45 seconds for 25 minutes. **b** *S. aureus* grown to late exponential phase was treated with the indicated compounds for 10 minutes prior to quantifying luminescence of the cell pellet and associated supernatant. Data represent the mean values from *n* = 3 biologically independent replicates ± SD. Statistical significance was determined by one-way ANOVA with Dunnett’s multiple comparisons test. The *p-*value of conditions with significance are indicated on the graph, otherwise comparisons were not significant (n.s.).

Next, we evaluated the membrane permeability by comparing intracellular versus extracellular ATP levels. Dual treated cells displayed increased extracellular ATP leakage (Fig. 3b) in a vancomycin-dose dependent manner. This data suggests that increased membrane permeability may result from the passive insertion of palmitoleic acid disturbing lipid packing, and disrupting the membrane barrier function^47^, rather than the formation of a structured transmembrane pore (Fig. 3a). This is also supported by previous work which has shown that unsaturated fatty acids increase membrane fluidity^46^.

### Vancomycin induces palmitoleic acid incorporation at the septum leading to disruption of functionalized fluid regions

Since dual treatment led to a gradual increase in membrane permeability, similar to effects previously seen with daptomycin which disrupts membrane organization and fluidity^30^. Recent studies show that the bactoprenol-bound precursors, such as lipid II, alter the phospholipid bilayer surrounding the bactoprenol molecule^48^. Moreover, to accommodate the long hydrocarbon tail of these molecules, a long-lived hydrophobic and fluid microdomain in the membrane, also known as regions of increased fluidity (RIFs), are generated^30,48^. By localizing bactoprenol-bound molecules in RIFs, these microdomains act as a “landing terrain” for lipid II binding antibiotics, as their targets can be distinguished in a ‘sea’ of other lipids^48^. Similar to the bulky hydrophobic nature of bactoprenol, the *cis* unsaturation in palmitoleic acid causes a ‘kink’ in its hydrocarbon tail and thus is more easily accommodated in a membrane environment with increased fluidity and hydrophobicity^49^. The specificity of palmitoleic acid synergy with vancomycin and bacitracin (Fig. 2) suggests that synergy depends on accumulation of bactoprenol-bound precursors. We hypothesized that vancomycin-induced accumulation of lipid II generates fluid hotspots for the insertion of palmitoleic acid which will ultimately disrupt cell envelope homeostasis, causing cell death. To investigate the effects of combination or monotherapy on *S. aureus* membrane fluidity and RIF organization we utilized high resolution fluorescence microscopy. The fluorescent lipophilic dyes nile red and DiI-C12 both insert into the membrane, preferentially accumulating in more fluid areas of the phospholipid bilayer resulting in increased fluorescence intensity^30,50,51^. Unlike *Bacillus subtilis, S. aureus* lacks a MreB cytoskeleton which is often associated with microscopically visible RIF organization^52^.

Although it has been suggested that *S. aureus* likely possess RIFs, the small cell size prevents observable RIFs in untreated cells stained with the RIF specific dye DiI-C12^42^. RIFs in *S. aureus* only become apparent when treated with membrane-active agents that induce larger RIF formation, as seen previously with rhodomyrtone and daptomycin^42,53^.

Consistent with previous studies, *S. aureus* control cells generally exhibit smooth, consistent fluorescence across the membrane using either nile red or DiI-C12 (Fig. 4a-c)^53,54^. Similarly, vancomycin monotherapy illustrated smooth membrane staining with either dye, although DiI-C12 fluorescence was brighter compared to control cells suggesting some overall increased fluidity in vancomycin treated cells (Fig. 4a-b). Palmitoleic acid also increased overall Dil-C12 florescense indicating some increased fluidity in the presence of the fatty acid alone (Fig. 4a-b). However, when cells were treated with both palmitoleic acid and vancomycin, large foci were easily observable using either nile red or Dil-C12, indicating the formation of large RIFs during dual-treatment (Fig. 4a-b, Supplementary Figure 2 and 3b). Even at sublethal concentrations where no cell death occurs, RIFs are still obvious in dual treated cells, though less pronounced (Supplementary Figure 3).

**Figure 4.**
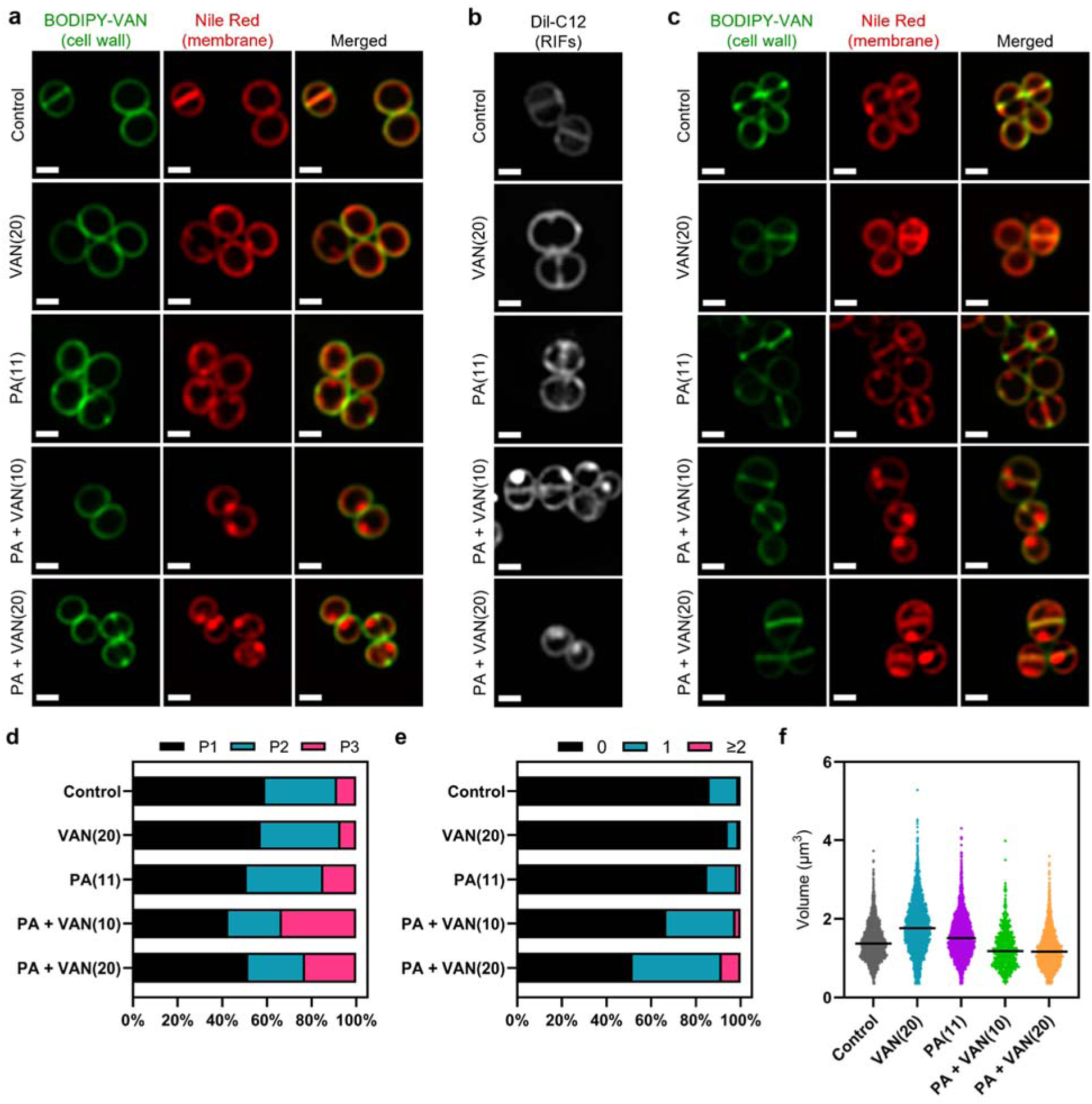
Dual treatment generates distinct fluid membrane patches. *S. aureus* RIFs visualized using fluorescent membrane dyes. *S. aureus* (HG003) treated with DMSO, PA (11 μg/ml), VAN (20 μg/ml), PA + VAN (10 μg/ml), or PA + VAN (20 μg/ml) for **a-b** 10 minutes or **c** 30 minutes. **a, c** HG003 was stained with fluorescent BODIPY labeled VAN (1 μg/mL) and nile red (5 μg/mL) for 5 minutes. **b** RIFs in HG003 were visualized by DiI-C12 (1 μg/ml). Cells were fixed prior to imaging on an agarose pad. Images are representative of the population. Scale bar, 1μm. **d-f** bioinformatic analysis of all cells treated for 30 minutes and imaged from 2 or 3 separate biological replicates. **d** the cell cycle was determined for each cell in a given treatment group and illustrated as a percent of the total population; Phase 1 (P1), phase 2 (P2), phase 3 (P3). **e** the number of nile red foci in a given cell was quantified for each treatment group and represented as a percent of the total population. **f** Scatter plot of cell size of each cell in the indicated treatment group. Black line represents the median. Statistical significance was determined by one-way ANOVA with Dunnett’s multiple comparisons test and all conditions had a p-value <0.0001, ****, compared to control.

The phenotypes visualized at 10 minutes were not transient, and images taken after 30 minutes displayed large foci only in dual treated conditions (Fig. 4c). Furthermore, dual treatment resulted in nearly 50% of cell population with at least one foci, often localized at the septum (Fig. 4e and Supplementary Fig. 4b). Interestingly, although there wasn’t a difference in cell size after 10 minutes (Supplementary Fig. 4c), dual treated cells had a significant decrease in cell volume after 30 minutes (Fig. 4f), suggesting that after prolonged exposure, dual treated treated cells shrink, likely reflecting the leakiness of the membrane. We also examined the impact on the cell cycle and found the distribution of the cell cycle in dual treated conditions shifted to a larger proportion of the population in phase 3 (Fig. 4d). Phase 3 is a short elongation step prior to an incredibly fast separation event^54,55^, and typically, only a small proportion of the cell population is found in phase 3, as observed in control and monotherapy-treated cells (Fig. 4d). These results illustrate that dual treatment generates large RIFs, leading to cell shrinkage and aberrant daughter cell splitting.

To further visualize membrane aberations in dual-treated cells, we performed transmission electron microscopy (TEM). We observed normal cellular structure and septal formation in control, palmitoleic acid alone, and vancomycin alone treated cells (Fig. 5a-c, e-g). In contrast, cells treated with both palmitoleic acid and vancomycin exhibited deformed septa that were misshaped and lacked an electron dense mid-line compared to control cells (Fig. 5d,h; yellow arrow). The dual treated cells additionally had an electron transparent region at the septa much thicker than the other conditions imaged and no discernable cell membrane or intermediate layer at the septum. Moreover, the dual treated cells had membrane invaginations that varied in size amongst the population of cells imaged (Fig. 5h, red arrow). Interestingly, these ultrastructural alterations of the cell membrane and septa parallel those reported for lipoglycopeptide treated *S. aureus*^56^.

**Figure 5.**
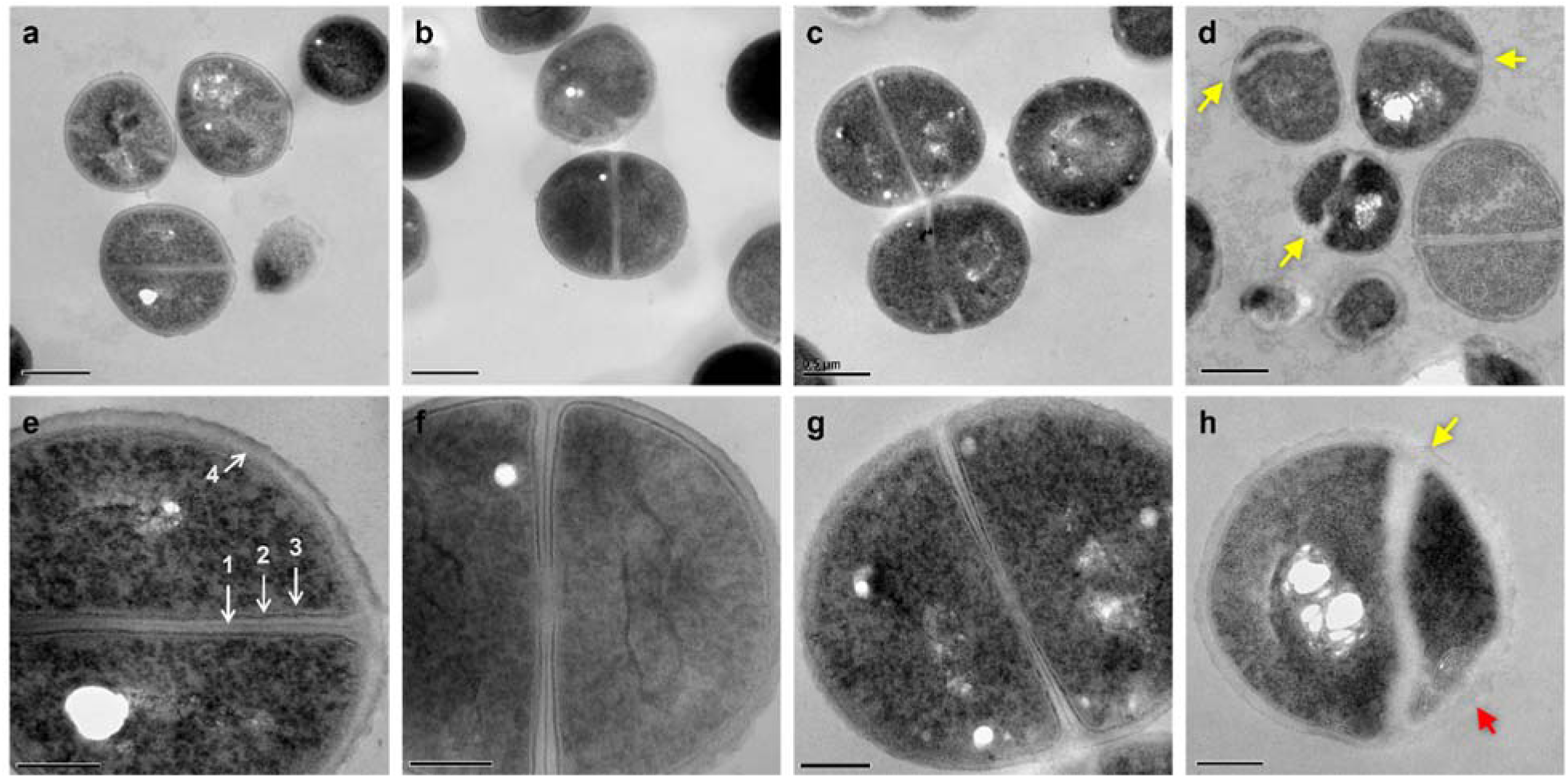
Dual treated cells cause septal aberrations. The ultrastructure of *S. aureus* cells treated for 30 minutes visualized by TEM. **a-h** *S. aureus* HG003 grown to mid-exponential phase was treated for 30 minutes prior to fixation with DMSO (a,e), 11 μg/ml PA (b,f), 20 μg/ml VAN (c,g) and PA + VAN (d,h). **a-c, e-g** micrographs of cells with a cross wall at mid-cell, while **d**,**h** show cells treated with PA+VAN have deformed septa (yellow arrows) and membrane invaginations (red arrows). Magnification of 50,000X with a 0.5μm scale bar (**a-d**) or magnification of 150,000X with a 200nm scale bar (**e-h**). (**e**) (1) electron dense midline of the septum, (2) electron dense intermediate layer, located between the cell membrane (3) and the cell wall (4).

### Dual treatment delocalizes cell division and peptidoglycan biosynthesis machinery

Disruption of RIF organization by other antibiotics have been found to displace or delocalize membrane-bound proteins, such as those of cell division and peptidoglycan biosynthesis^30,53^. The divisome of *S. aureus* is a multi-protein complex that forms a ring at the site of cell division^57^. Cell division requires the highly coordinated efforts of peptidoglycan synthesis, hydrolysis, and turgor pressure^54,58^. EzrA is an essential divisome protein that acts as a scaffold and regulator for the proper formation of the division ring at the septum^57,59^. To determine whether the septal defects visualized with TEM are caused by an improper localization of the divisome complex, we utilized a *S. aureus* strain with an EzrA-GFP fusion^60^. We found that the control and vancomycin treated cells maintained EzrA fluorescence at the septum as a single ring at the center of the cell (Fig. 6a). In contrast, we observed two distinct phenotypes following dual treatment with palmitoleic acid and vancomycin: delocalization of EzrA throughout the cell or multiple distinct rings within a single cell (Fig. 6a). Interestingly, we observed a small subpopulation of cells treated with palmitoleic acid monotherapy with delocalized EzrA (Fig. 6a). This is consistent with a previous study that suggested low concentrations of antimicrobial lipids, including palmitoleic acid, exert bacteriostatic activity as a result of momentary cell division defects prior to cell recovery^31,61^.

**Figure 6.**
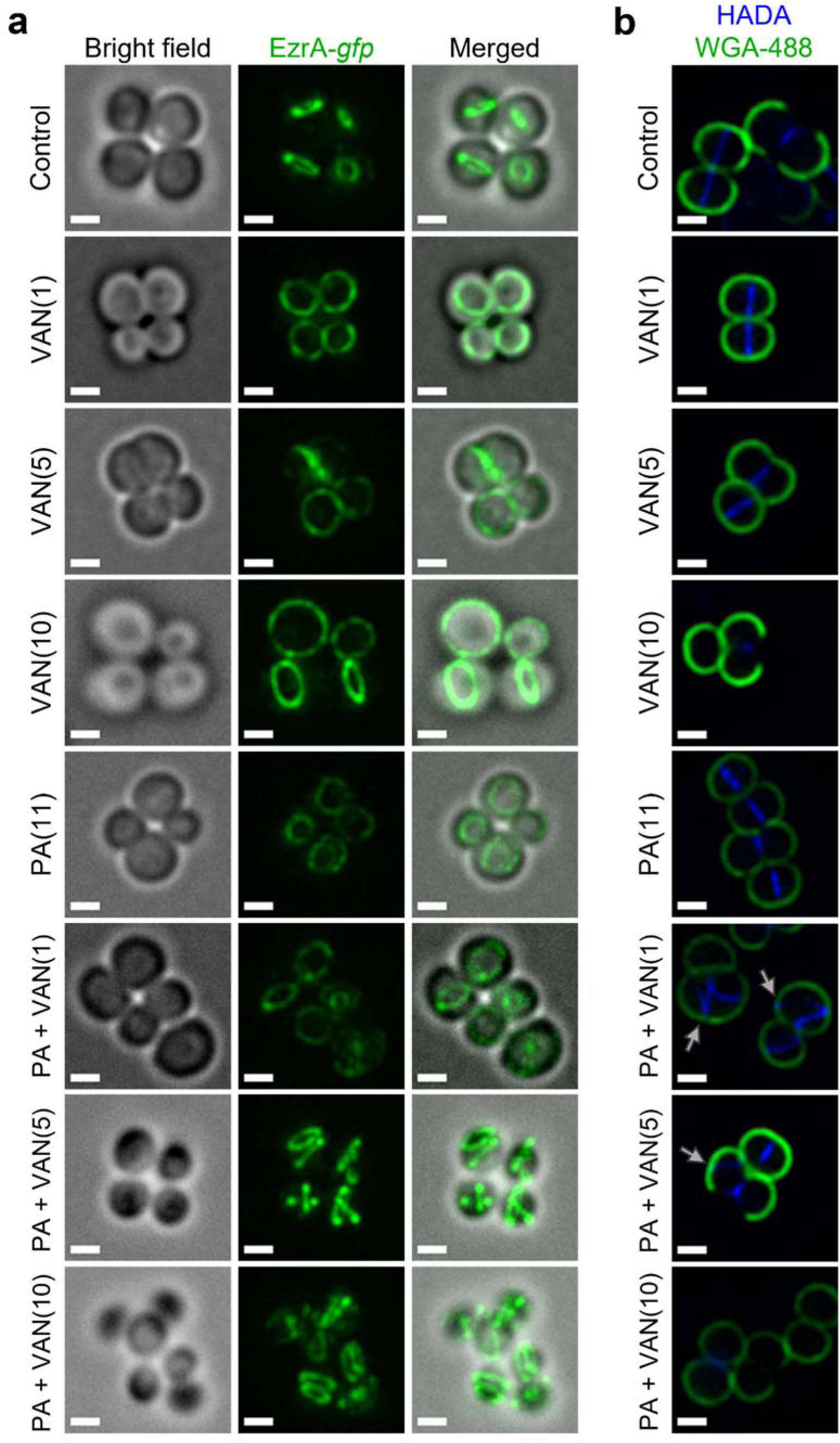
Dual treatment causes protein delocalization. Localization of cell division and peptidoglycan biosynthesis machinery after 10 minutes. **a** HG003 with chromosomal *ezrA-gfp* was grown to exponential phase prior to treatment with DMSO (control), VAN (1, 5, 10, or 20 μg/ml), PA (11 μg/ml), or PA + VAN at indicated concentrations in parentheses. **b** HG003 grown to exponential phase and treated for a total of 10 minutes as indicated, HADA (blue, 250 μM) was added after 5 minutes. The compounds and HADA dye were washed from the cells and stained with WGA (green, 1 μg/ml) for 5 minutes and then fixed. Gray arrows illustrated aberrant localization of peptidoglycan synthesis. **a-b** After treatment, cells were fixed prior to imaging on an agarose pad. Scale bars are 1μm.

Since peptidoglycan synthesis is directly linked to cell division, we next aimed to determine if dual treatment alters peptidoglycan synthesis using HADA, a fluorescent D-amino acid that is incorporated into newly synthesized cell wall, and wheat germ agglutinin (WGA) that stains mature cell wall^62^. After 10 minutes, the lower vancomycin monotherapy concentrations had yet to inhibit cell wall biosynthesis, but the highest concentration had shut down peptidoglycan biosynthesis in a majority of the cells in any given field (Fig. 6b). In all control or monotherapy groups, HADA was correctly located at the septa (Fig. 6b). In contrast, when palmitoleic acid was combined with lower vancomycin concentrations that allowed HADA incorporation, we saw aberrant peptidoglycan incorporation (Fig. 6b, gray arrows). Peptidoglycan biosynthesis is a highly regulated process that occurs at the site of cell division prior to rapid daughter cell separation^58^. Dual treated cells instead began to initiate peptidoglycan synthesis at additional division planes prior to daughter cell separation further indicating cell division or cytokinesis defects (Fig. 6b, gray arrows)^63^. Together, these data suggest that vancomycin in combination with palmitoleic acid promotes divisome and peptidoglycan biosynthesis delocalization, leading to lethal aberant septa formation.

### Palmitoleic acid sensitizes antibiotic resistant isolates to vancomycin killing

One strategy to reduce the rise in resistance while also sensitizing already resistant and tolerant populations is to utilize chemotherapies with multiple bacterial targets^13^. Because the combination of palmitoleic acid and vancomycin targets multiple essential processes, we aimed to determine if dual treatment can re-sensitize resistant isolates to vancomycin. Despite the decreased susceptibility of characterized VRSA and VISA clinical isolates to vancomycin^64^, dual treatment resulted in similar killing kinetics to MSSA, killing 99.9% of the cells after only 30 minutes (Fig. 7a,b and 1b). Additionally, palmitoleic acid potentiated vancomycin killing of other Gram-positive bacteria, including *Enterococcus faecalis, Bacillus subtills, Enterococcus faecium*, and *Staphylococcus epidermidis* (Fig. 7c-g). Importantly, palmitoleic acid was able to potentiate vancomycin killing of VRE (Fig. 7d). However, palmitoleic acid did not potentiate killing of the Gram-negative bacterial species *E. coli* (Fig. 7h), likely due to the outer membrane. These data illustrate the potency of palmitoleic acid as a vancomycin adjuvant against various Gram-positive bacteria, including those exhibiting vancomycin resistance.

**Figure 7.**
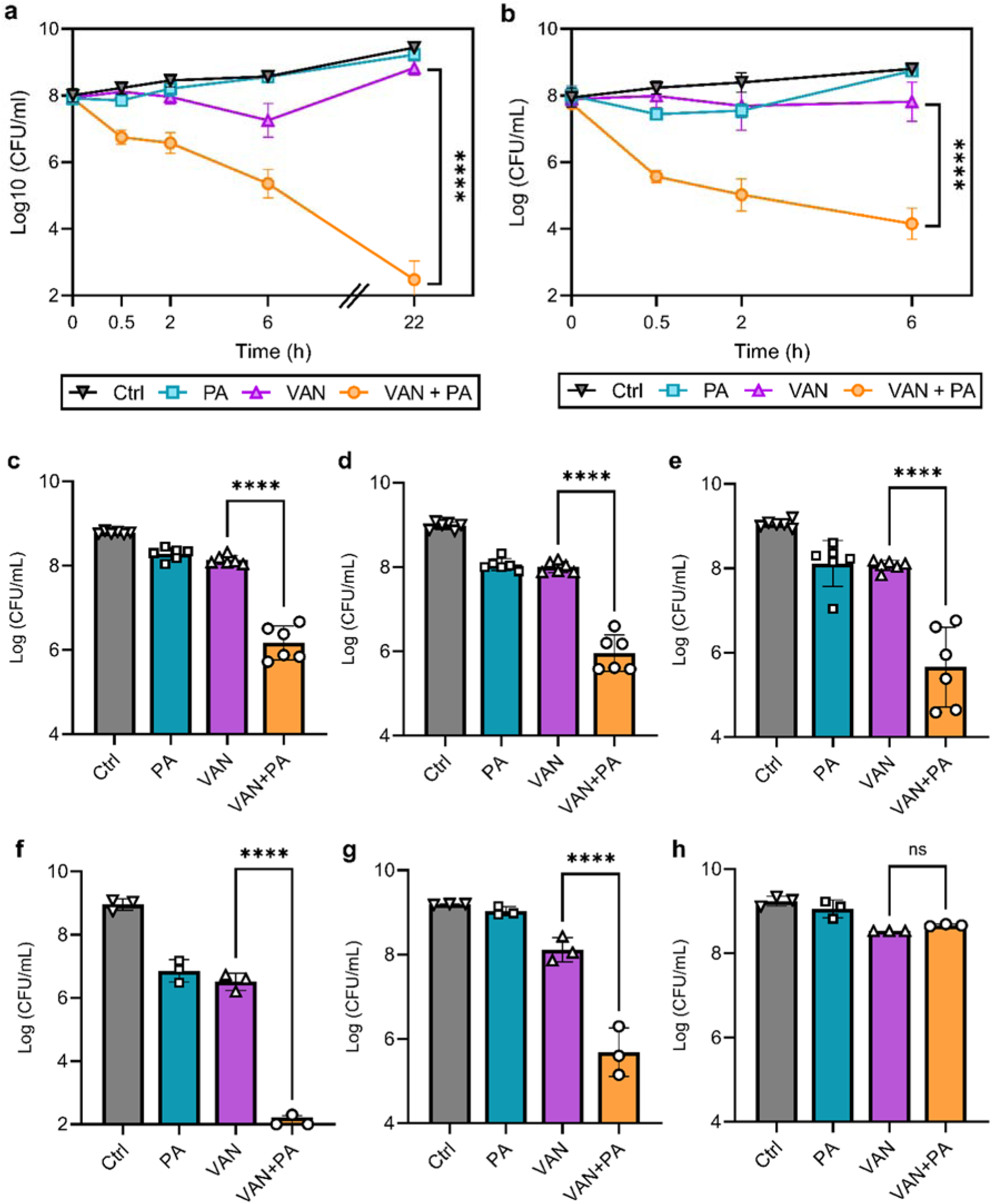
Palmitoleic acid re-sensitizes vancomycin resistant isolates. Combination therapy leads to rapid killing of Gram-positive bacterial species. **a** VISA, and **b** VRSA strains were grown to exponential phase prior challenge with DMSO (Ctrl), PA (11 μg/ml), VAN (20 μg/ml), or VAN + PA. At the indicated time points an aliquot was removed, washed, and plated to enumerate survivors. PA potentiation of other Gram-positive bacteria was determined using **c** *Enterococcus faecalis* strain OG1, **d** vancomycin-resistant *E. faecalis* strain V583, **e** *Enterococcus faecium*, **f** *Bacillus subtilis*, **g** *Staphylococcus epidermidis* CSF41498, and Gramnegative bacterial species **h** *Escherichia coli* MG1655. Strains were grown to exponential phase prior to exposure to DMSO, PA, VAN, or PA + VAN (*see Supplementary Table 2 for concentrations)*. After 2h, an aliquot was removed, washed, and plated for CFU enumeration. Data represents *n* = 6 or 3 biologically independent replicates with error bars as SD. Significance was evaluated using students two-tailed unpaired *t*-test between VAN and VAN + PA conditions.

## Discussion

Although new therapeutics are in the pipeline, preservation and improvement of pre-existing therapies is important for combatting rising rates of antibiotic treatment failure and antibiotic resistance. Vancomycin is a gold standard antimicrobial as it took nearly 40 years after its introduction to the clinic before the first resistant isolate appeared^65^. Unfortunately, vancomycin resistance is becoming more and more prevalent, particularly amongst Enterococci where over half of all *E. faecium* isolates now exhibit vancomycin resistance^7^. Additionally, under certain environmental conditions including high bacterial density, vancomycin is poorly bactericidal, facilitating antibiotic tolerance, infection recurrence, long hospital stays, and prolonged courses of parenteral antibiotics^23,24,66^. Here, we’ve identified a vancomycin adjuvant, palmitoleic acid, that potentiates vancomycin killing against multiple Gram-positive bacteria. Importantly, palmitoleic acid increased the rate of killing of vancomycin at high bacterial densities, killing over 99.9% of the bacterial populations of most isolates within 30 minutes, improving significant pitfalls of vancomycin monotherapy.

Research focused on antimicrobisl activity of UFAs dates back nearly a century due to their potency at high concentrations and their lack of toxicity to mammalian tissue^67-70^. UFAs are found in nasal secretions and on the skin as an important natural protective barrier, in addition to being commonly found in our diet^34,61,71^. Mice deficient in skin monounsaturated fatty acid production show increased susceptibility to *S. aureus* in a cSSTI infection model^72^. Despite immense interest in antimicrobial fatty acids over the decades, a concise mechanism of action ascribed to UFAs remains unclear. Some effects attributed to UFA antibacterial activity include disruption of signal transduction, membrane depolarization, disruption in energy production, inhibition of fatty acid synthesis, increased permeability, and membrane dissolution^25,43,44,73,74^. We find that although dual treatment seems to increase membrane permeability, it is not via pore formation but rather a localized increase in membrane fluidity that prevents proper barrier function. However, increased permeability is unlikely to be the main cause of cell death, as most CMAAs tested have been shown to also increase membrane permeability, but did not potentiate vancomycin killing^18,19,31,32^. Together these results suggest that an ideal vancomycin adjuvant, such as palmitoleic acid, must meet specific criteria to cause the necessary membrane perturbations that potentiate vancomycin activity.

Interestingly, previous work has found that accumulation of lipid II is saturated after 30 minutes of vancomycin treatment^5^. We speculate that the rapid killing seen by 30 minutes when vancomycin is combined with palmitoleic acid relies on maximizing lipid II accumulation on the cell surface to create the ideal insertion point for palmitoleic acid^5,48^. The insertion of palmitoleic acid into the ideal microenvironment around lipid II increases the disordered phospholipid environment surrounding the peptidoglycan monomer, subsequently creating large RIFs that we visualized with nile red and DiI-C12 staining. Furthermore, we suggest accumulation of lipid II and insertion of palmitoleic acid are two actions that occur in parallel and are likely the central events that must occur to mediate synergy between these compound.

Lipid II is not only a simple building block of peptidoglycan, but also has numerous crucial roles in the orchestration of essential processes in bacteria. In fact, lipid II appears to be the molecular signal for fine-tuning cell envelope biosynthesis, including during cell division^75^.

Interfering with lipid II localization subsequently disrupts the highly coordinated cell wall biosynthesis and divisome machinery causing complex downstream effects that, when combined with membrane damage, are irreparable^4,76^. We visualized these downstream effects as defective septal formation in TEM and multiple EzrA rings. Our results suggest that lipid II sequestration is significantly perturbed in the dual treatment.

The capacity of CMAAs to sensitize vancomycin resistant isolates is not without precedence. Interestingly, previous studies have reported that prolonged treatment with antimicrobial peptides re-sensitized VRSA to vancomycin^77,78^. Here, we find that administration of palmitoleic acid simultaneously with vancomycin re-sensitizes resistant strains to vancomycin killing. The most common mechanism of vancomycin resistance is maintenance of a plasmid containing the *vanA* operon. Sensing of vancomycin activates transcription of the *vanA* operon and generates an altered form of lipid II that has a terminal D-Ala-D-Lac on the stem peptide which cannot be bound by vancomycin^8^. Although it is unclear how palmitoleic acid re-sensitizes resistant isolates to vancomycin, it is possible that the rapid bactericidal activity of the combination leaves little time for VRSA or VRE isolates to induce expression of the *van* genes in response to vancomycin.

Because palmitoleic acid is already a natural protectant on the skin surface and has been shown accelerate wound healing in a rat model when applied topically^79^, palmitoleic acid could potentially be utilized as a topical therapeutic in conjunction with intravenous vancomycin for cSSSIs. Topical ointments are preferred as they provide localized high concentrations of antimicrobials without the risk of systemic toxicity. The leading over-the-counter topical ointment, Polysporin, contains high concentrations of bacitracin for acute skin infections^80^.

Unfortunately, resistance to bacitracin is widespread in Staphylococci and Enterococci diminishing its utility against these infections^81,82^. Since we find that palmitoleic acid potentiates bacitracin killing, it is possible that the addition of palmitoleic acid to these common ointments could improve their efficacy. However additional work is needed to evaluate the potential therapeutic efficacy of palmitoleic acid as a topical ointment in vivo.

In summary, the findings presented here demonstrate a promising approach to enhancing approved therapeutics that bypasses the need to identify novel antimicrobial compounds. Focusing on antibiotic adjuvants that are naturally non-toxic to improve the efficacy of essential antibiotics can combat antimicrobial resistance and tolerance. While unraveling the mechanism of action of lipid II and membrane targeting antimicrobials has consistently been challenging, we provide an important and more detailed piece of the puzzle on how targeting lipid II and the membrane in general leads to potent synergy.

## Materials and Methods

### Bacterial growth and conditions

*S. aureus* strains HG003 (MSSA)^83^, HG003 *ezrA-gfp*, LAC (CA-MRSA)^84^, SA770 (VISA), and VRS2 (VRSA)^64^, *B. subtilis* strain RT275, *E. coli* strain MG1655, and *S. epidermidis* strain CSF41498 were routinely cultured in tryptic soy broth (TSB, Remel) at 37°C with 225rpm shaking. *E. faecalis* strains OG1 and V583, and a clinical *E. faecium* isolate^85^ were cultured in Brain Heart Infusion (BHI) Broth (Oxoid) at 37°C with shaking (*E. faecium*) or statically (*E. faecalis* strains). The EzrA-GFP was transduced from the chromosome of ColpSGEzrA-GFP^58^ to HG003 by phage transduction using phage 80α as previously described^86^.

### In vitro antibiotic survival assays

Antibiotic survival assays were performed as previously described^36^. Briefly, overnight cultures were diluted 1:1000 into fresh media and grown to ∼2×10^8^ CFU/ml aerobically at 37°C, 225rpm (unless otherwise indicated) prior to antibiotic challenge as indicated in the figure legends. Concentrations of antibiotics used in survival assays can be found in Supplementary table 2. At the indicated time points, an aliquot of cells was washed twice with PBS, serially diluted, and plated on tryptic soy agar (TSA) to enumerate CFU. Statistical analysis of vancomycin synergy with CMAAs after 6 hours was performed by ordinary one-way ANOVA with Dunnett’s multiple comparisons, comparing the mean of each CMAA+VAN combination with the mean of the VAN control. Statistical significance of antibiotic survival assays displaying multiple time points was determined by using a two-tailed unpaired Student’s *t*-test with a 95% confidence interval using the values of the final time point.

### Membrane integrity assays

Membrane potential was measured using the fluorescent dye 3,3′-dipropylthiadicarbocyanine iodide (DiSC_3_(5), Invitrogen) as previously described^45^. Briefly, measurements were carried out in a Synergy H1 (BioTek) plate reader equipped with 610nm excitation and 660nm emission filters. *S. aureus* HG003 was grown in TSB as described above. After reaching the desired starting CFU, 135μL of culture was added to a black flat bottom 96-well polystyrene plate that was non-binding (Corning, 3991) to prevent the necessity of BSA being added. DiSC_3_(5) was added to each well at a final concentration of 7μM. The optimal DiSC_3_(5) concentration was determined by evaluating the difference between positive (gramicidin) and negative (DMSO) controls at various DiSC_3_(5) concentrations. The maximum dynamic range between controls at the cell density used was determined to be 7μM. The dye was allowed to equilibrate for 3 minutes prior to adding the compounds of interest. Gramicidin (1uM) was used as a positive control. Reads were taken every 45 seconds for 25 minutes while shaking.

ATP levels were evaluated using the BactiterGLO kit (Promega) per manufacturer’s specifications and normalized to protein content in the cell pellet as determined using a Bradford assay.

### Fluorescence microscopy

All imaging was performed using *S. aureus* HG003 grown as described above prior to challenge with compounds at concentrations and time points indicated in figure legends.

To evaluate morphological changes, 1ml of culture was incubated for 5 minutes at 37°C with 5 μg/ml of nile red (Sigma) to stain the membrane and the cell wall was labeled with 1 μg/ml of an equal mixture of vancomycin (Alfa Aesar) and a BODIPY FL conjugate of vancomycin (Invitrogen). Excess dye was removed by washing with TSB. To determine if membrane perturbations were RIFs the lipophilic dye DiI-C12 (Invitrogen) was used as described^50^. Briefly, overnight cultures were diluted 1:1000 into fresh TSB containing 1% DMSO and 1 μg/ml DiI-C12. Cells were grown to ∼2×10^8^ CFU/ml and washed four times with pre-warmed TSB containing 1% DMSO. Cells were then treated for 10minutes and washed to remove excess antibiotic and dye. Washes and incubation steps were performed in a warm room to maintain samples at 37°C for optimal DiI-C12 solubility. For visualization of divisome proteins, overnight cultures of HG003 *ezrA-gfp* were diluted 1:1000 into fresh TSB and grown to ∼2×10^8^ CFU/ml. Cells were then treated for 10 minutes prior to washing with TSB. To visualize peptidoglycan biosynthesis, 1ml of culture was incubated with the fluorescent D-amino acid HADA (Tocris Bioscience) at a final concentration of 250μM for 5 minutes. The cells were subsequently washed with TSB and incubated with WGA tagged with Alexa Fluor 488 (WGA-488, Invitrogen) at a final concentration of 2 μg/ml for 5 minutes; excess WGA-448 was removed by washing with TSB. Both incubation steps were carried out at 37°C with agitation.

After staining, *S. aureus* was fixed with 4% paraformaldehyde made fresh day of for 20 minutes at room temperature with agitation. Cells were then washed with PBS and resuspended in 1/10^th^ the starting volume. Samples were imaged within 24hrs of fixation by pipetting 2μl of sample onto an agarose pad (made as previously described^87^) and mounted onto a #1.5 cover slip. Image Z-stacks were acquired using an Olympus IX81 inverted microscope fitted with a Hamamatsu ORCA-Flash4.0 V3 camera, X-Cite XYLIS XT720L (385) illumination, 100X/1.4 Oil UPlan S Apo PSF objective lens, and Metamorph 7.10 acquisition software (Molecular Devices). Z spacing was 0.2 μm, and pixel size in XY was 0.064 μm. Our Z-stacks ranged in thickness from 2.6-4.0 μm. We acquired images for all channels at each Z position before moving to the next Z position. Nile red was imaged with 560/25 excitation and 607/36 emission filters. BODIPY-VAN, EzrA and WGA-488 were imaged with 485/20 excitation and 525/30 emission filters. HADA was imaged with 387/11 excitation and 440/40 emission filters. DiI-C12 was imaged with 572/35 excitation and 632/60 emission filters. All images were acquired with the same exposure settings. No pixels had saturated intensities. Z-stacks were subsequently deconvolved using Autoquant software version 3.1.3 with default settings. All images in a figure were processed in FIJI and have the same display adjustments. A single Z-plane that was determined to be the middle cross-section of *S aureus* cells in the field of view was used for representative images of nile red, VAN-FL, WGA-488, and DiI-C12. A max projection of all Z-stacks was used for the representative images of HADA and EzrA-GFP.

### Image analysis

Image analysis was performed using Python with scikit-image (0.18.2) image processing library^88^ and Fiji (v1.53q)^89^. The full analysis code is available on Github (https://github.com/fjorka/bacteria_pa_van_analysis) while all raw imaging data and key processed steps were deposited in Zenodo repository.

For quantification of bacteria cell features (volume, phase and foci number), a middle crosssection from confocal z-stacks was used. This plane was selected automatically by finding the frame with maximum cell area from thresholded (Otsu) images of the cell wall (BODIPY-VAN). Detection of cells was performed in the membrane channel (Nile Red) using cellpose segmentation algorithm^90^. Total number of detected cells (all conditions) >32k. Volume of cells was calculated based on the major and minor axis length of segmented cells as described previously^54^. Membrane foci were segmented using Maximum Entropy Threshold (Fiji). Phases of the cell cycle were determined using a classification network trained using fastai deep learning library (2.5.3)^91^. Briefly, a Resnet50 pre-trained network (fastai) was trained using manually annotated sets of cells (phase 1 - 150, phase 2 - 150, phase 3 - 150, missegmented cells - 80) achieving error rate <4%. Trained classifier was used to predict the cell cycle phase of remaining cells.

### Transmission Electron Microscopy

*S. aureus* was grown and challenged as described above. Cultures were centrifuged at 1500 x *g* and the supernatant was removed. Bacterial cells pellets were resuspended in 4% paraformaldehyde/2.5% glutaraldehyde in 0.15M sodium phosphate buffer, pH 7.4, for 1hr at room temperature and stored at 4°C. Following 3 washes with 0.15M sodium phosphate buffer, pH 7.4, the fixed cell pellets were post-fixed in 1% osmium tetroxide in 0.15M sodium phosphate buffer, pH 7.4 for 45 minutes and washed 3 times with deionized water. The samples were gradually dehydrated with ethanol (30%, 50%, 75%, 100%, 100%), and propylene oxide. The cell pellets were infiltrated with a 1:1 mixture of propylene oxide for and Polybed 812 epoxy resin overnight, followed by a 1:2 mixture of propylene oxide for and Polybed 812 epoxy resin for 3 hours and transferred to 100% resin to embed overnight (Polysciences, Inc.,Warrington, PA). Ultrathin sections (70-80 nm) were cut with a diamond knife and mounted on 200 mesh copper grids followed by staining with 4% aqueous uranyl acetate for 12 minutes and Reynold’s lead citrate for 8 minutes^92^. Samples were observed using a JEOL JEM-1230 *transmission* electron microscope operating at 80kV (JEOL USA, Inc., Peabody, MA) and images were acquired with a Gatan Orius SC1000 CCD Digital Camera and Gatan Microscopy Suite 3.0 software (Gatan, Inc., Pleasanton, CA).

## ACKNOWLEDGEMENTS

This work was supported in part by NIH grants R01AI137273, Burrough’s Wellcome Fund, and Cystic Fibrosis Foundation Research Grant to B.P.C. We thank Marianna Pinho for providing COL-*ezra-gfp*, Kim Lewis for providing VISA and VRSA isolates, and Jeremiah Faith for providing the clinical *E. faecium* isolate. We thank Pablo Ariel at the Microscopy Service Laboratory (MSL) at UNC for helping optimize fluorescence microscopy experiments. We thank Kristen White at the MSL for performing TEM imaging and processing. The MSL, Department of Pathology and Laboratory Medicine, is supported in part by P30 CA016086 Cancer Center Core Support Grant to the UNC Lineberger Comprehensive Cancer Center. The image analysis was performed at the Bioinformatics and Analytics Research Collaborative (BARC) which is supported in part by the UNC School of Medicine Strategic Plan and Office of Research. KMK is supported by grant number 2020-225716 from the Chan Zuckerberg Initiative DAF, an advised fund of Silicon Valley Community Foundation. We thank Virginie Papadopoulou, Phil Durham, Lauren Radlinski, Baggio Evangelista, and Joey Ragusa for thoughtful discussions.

## COMPETING INTEREST STATEMENT

B.P.C and S.E.R are co-inventors on a provisional patent describing the use of membrane acting agents for potentiating antibiotic efficacy. A.E.S, K.M.K, J.E.B, J.B.P., and D.T.B declare no competing interests.

