## Supplemental Figures for "Antibiotic-induced accumulation of lipid II sensitizes bacteria to antimicrobial fatty acids"

**Supplementary figures**

| Supplementary Table 1. Structures of cell-membrane acting agents | | |
|---|---|---|
| 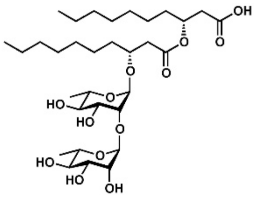 Rhamnolipids (Biosurfactant) | 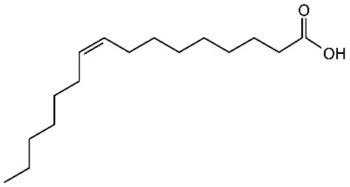 Palmitoleic acid (UFA, C16:1) | 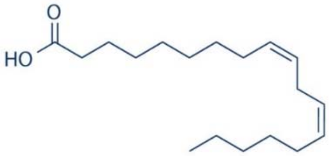 Linoleic acid (UFA, C18:2) |
| 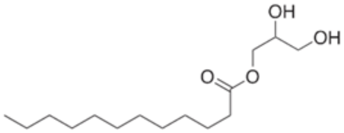 Glycerol Monolaurate (Monoglyceride) | 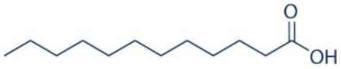 Lauric acid (Saturated fatty acid, C12:0) | |
| 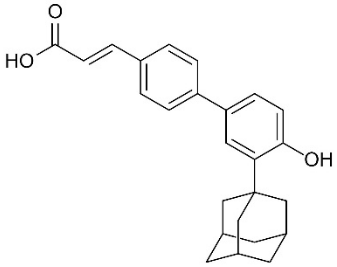 Adarotene (Synthetic retinoid) | 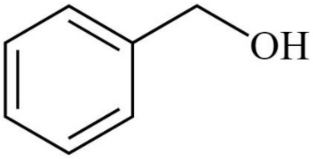 Benzyl Alcohol (General membrane fluidizer) | |

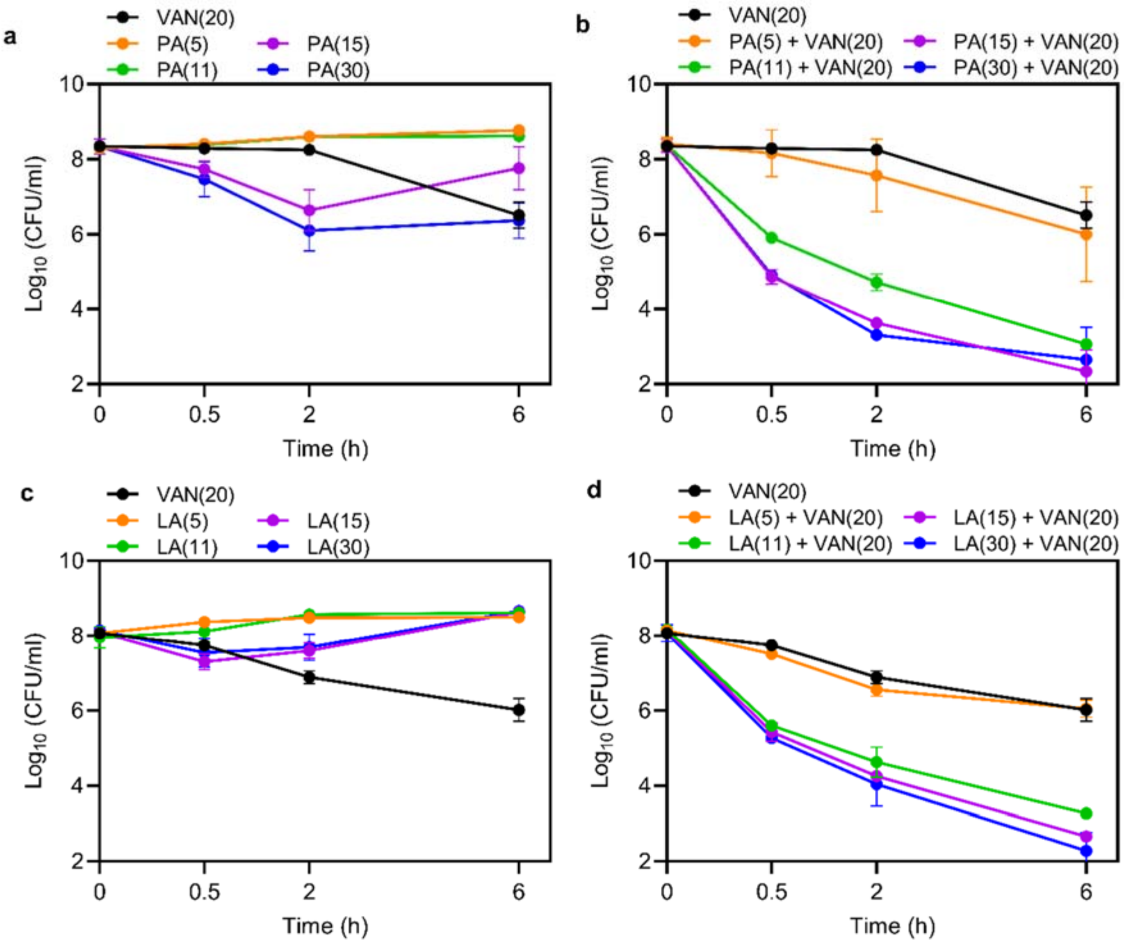


**Supplementary Figure 1. Identification of sublethal concentration of UFAs.** HG003 was grown to exponential phase prior to challenge with 5, 11, 15, or 30 µg/ml of **a** PA alone, **b** PA in combination with VAN (20 µg/ml), **c** LA alone, or **d** LA in combination with VAN. An aliquot of cells was removed at the indicated time point, washed, and plated for CFU. Data represent the mean values from *n* = 3 biologically independent replicates ± SD.

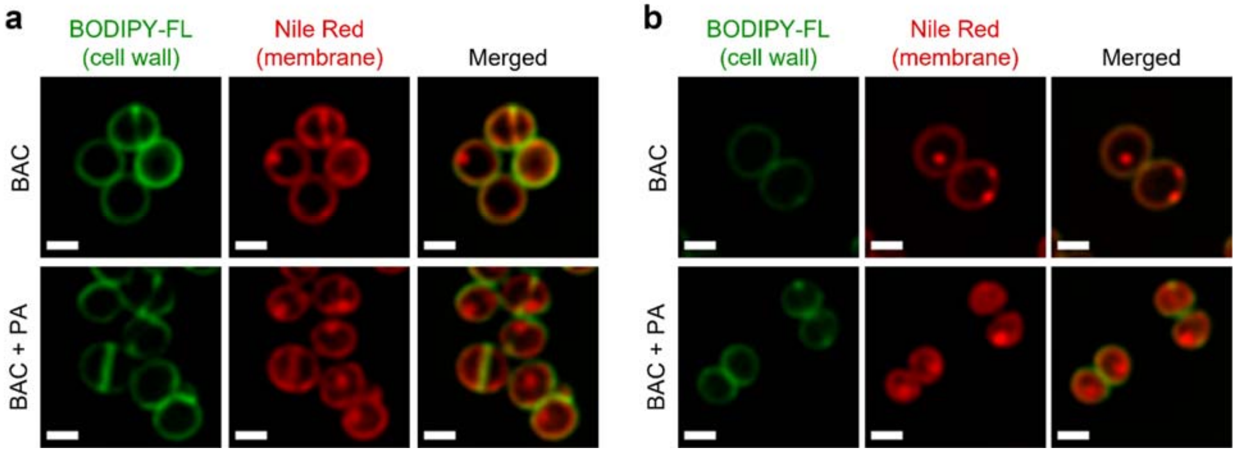


**Supplementary Figure 2. PA combined with bacitracin causes distinct fluid membrane patches.** Fluid foci microdomains in the membrane visualized using nile red. *S. aureus* HG003 treated with DMSO, PA (11 µg/ml), bacitracin (BAC, 250 µg/ml), or PA + BAC for **a** 10 minutes or **b** 30 minutes. Cells were stained with BODIPY labeled VAN (1 µg/ml) and nile red (5 µg/ml) for 5 minutes. Cells were fixed prior to imaging on an agarose pad. Images are representative of the population. Scale bar, 1µm.

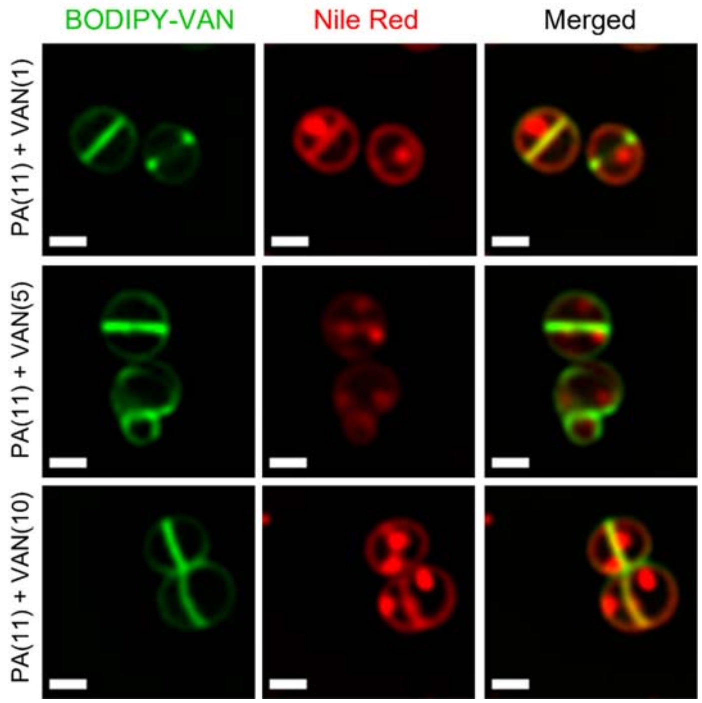


**Supplementary Figure 3. Sublethal concentrations of dual treatment still induce membrane aberrations.** *S. aureus* RIFs visualized using nile red. HG003 treated with DMSO, PA (11 μg/ml), or PA + VAN (1, 5, or 10 μg/ml) for 10 minutes. HG003 was stained with fluorescent BODIPY labeled VAN (1 μg/mL) and nile red (5 μg/mL) for 5 minutes. Cells were fixed prior to imaging on an agarose pad. Images are representative of the population. Scale bar, 1μm

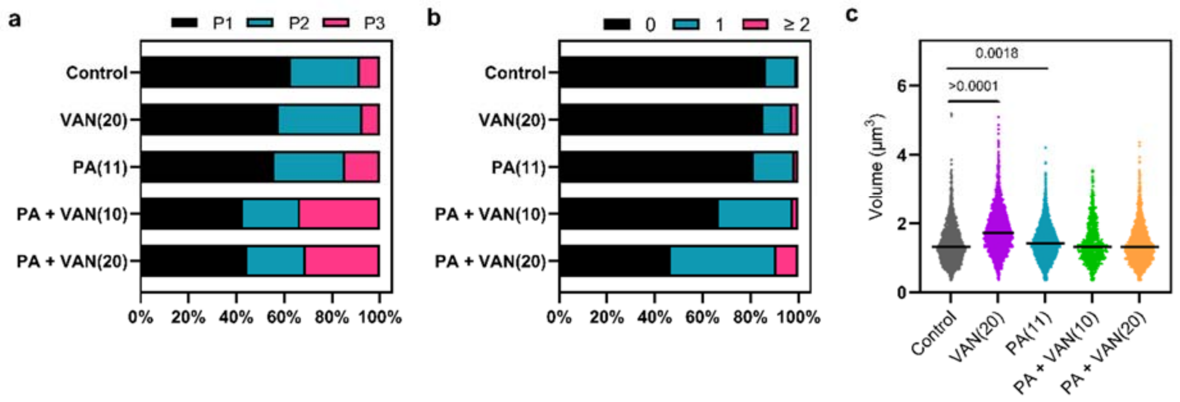


**Supplementary figure 4. Quantification of images taken after 10 minutes of antibiotic exposure.** Bioinformatic analysis of all cells treated for 30 minutes and imaged from 2 or 3 separate biological replicates. **a** the cell cycle was determined for each cell in a given treatment group and illustrated as a percent of the total population; phase 1 (P1), phase 2 (P2), phase 3 (P3). **b** the number of nile red foci in a given cell was quantified for each treatment group and represented as a percent of the total population. **c** Scatter plot of cell size of each cell in the indicated treatment group. Black line represents the median. The *p*-value of conditions with significance are indicated on the graph, otherwise comparisons were not significant (n.s.).

**Supplementary Table 2. Antimicrobial concentrations used in antibiotic susceptibility assay by strain**

| Strain | Vancomycin (µg/ml) | Palmitoleic acid (µg/ml) |
|---|---|---|
| *E. faecalis* OG1 | 50 | 30 |
| *E. faecalis* V583 | 50 | 30 |
| *E. faecium* | 50 | 11 |
| *S. epidermidis* CSF41498 | 50 | 11 |
| *E. coli* MG1655 | 50 | 50 |
| *B. subtilis* | 20 | 11 |
| *S. aureus* | 20 | 11 |
